# Parafoveal words can modulate sentence meaning: Electrophysiological evidence from an RSVP-with-flanker task

**DOI:** 10.1101/2021.07.06.451256

**Authors:** Nan Li, Olaf Dimigen, Werner Sommer, Suiping Wang

## Abstract

During natural reading, readers also take up some visual information from not-yet-fixated words to the right of the current fixation and it is well-established that this parafoveal preview facilitates the subsequent foveal processing of the word. However, the extraction and integration of word meaning from the parafoveal word and its possible influence on the semantic sentence context are controversial. In the current study, we recorded event-related potentials (ERPs) in the RSVP-with-flankers paradigm to test whether and how updates of sentential meaning that are based only on parafoveal information influence the subsequent foveal processing. Using Chinese sentences, the sentence congruency of parafoveal and foveal target words were orthogonally manipulated. In contrast to previous research, we also controlled for potentially confounding effects of parafoveal-to-foveal repetition priming (identity preview effects) on the N400. Crucially, we found that the classic effect of foveal congruency on the N400 component only appeared when the word in preview had been congruent with sentence meaning; in contrast, there was no N400 when the preview word had been incongruent. These results indicate that sentence meaning rapidly adapts to parafoveal preview, which already changes the context for the then fixated word. We also show that a correct parafoveal preview generally attenuates the N400 once a word is fixated, regardless of congruency. Taken together, our findings underline the highly generative and adaptive framework of language comprehension.

---

The reading of sentences differs in many regards from the reading of single words. In particular, readers of sentences actively generate internal, message-level representations that are incrementally updated with the unfolding sentence input. For example, the contextually appropriate meaning of an ambiguous word is facilitated by an appropriate context (e.g. Swinney, 1979), syntactic parsing is constrained by knowledge about the sentence (e.g. Taraban & McClelland 1988), and textual context can tear down the local congruency of the sentence (e.g. Albrecht & O’Brien, 1993). Findings like these underline the generative and highly adaptive nature of language comprehension (Wittrock, 1974, 1989).

During reading, processing of a word usually starts while it is still located in the parafoveal region of the visual field. However, due to the sharp drop-off of visual acuity from fovea (central 1-2°) to the parafovea (eccentricity of 2-5°) and periphery, the amount of information gleaned from not-yet-fixated words is limited (Rayner, 1998). Nevertheless, information from parafoveal preview clearly facilitates reading (McConkie & Rayner, 1975; Rayner, 1975). This raises the question whether readers generate contextual information based on parafoveal words. Put differently, is the sentence context updated not only by the currently fixated word, but already on the basis of parafoveal information? And if so, is the context generated by parafoveal words qualitatively similar to that generated by fixated foveal words? In the current study, we investigated whether readers can use parafoveal information to adjust the sentence meaning, thereby already changing the context for the fixated word.

Rather than passively consuming incoming information, generative theories view the neurocognitive system as actively constructing relations between parts of the text and drawing inferences from them (Wittrock, 1974). For example, the Bayesian Reader theorem predicts that readers make use of any source of available information to optimally recognize words (Delaney-Busch, Morgan, Lau, & Kuperberg, 2019; Kuperberg, 2016; Norris, 2006). Sentence meaning generated through parafoveal preview can be considered as one such source of information. If so, sentence processing should adapt to words in the parafoveal preview in a similar way as to words that are directly fixated. Conversely, however, the usefulness of parafoveal information is limited by low acuity and crowding, which degrades the input from the parafoveal visual field (Bouma, 1970; Pelli & Tillman, 2008; Rayner 1998). For example, Pernet, Uusvuori, and Salmelin (2007) found that priming effects by parafoveal prime words were weaker than for foveal primes and measurable only at short prime-target intervals. It therefore remains to be seen whether and how information gleaned from parafoveal words affects sentence meaning.

In the current study, we tested whether parafoveal word meaning modulates sentence meaning and thereby changes the sentential context for the subsequent direct fixation on the word. To this aim, we orthogonally manipulated whether the parafoveal word and the subsequently fixated foveal word were congruent with the sentence context or not (e.g. Barber, van der Meij, & Kutas, 2013; Li, Niefind, Wang, Sommer & Dimigen, 2015). In addition, to control for confounding effects of repetition priming (identity preview benefit) on foveal N400 amplitude, we also manipulated the lexical identity between the previewed word and the foveal word. A modulation of the N400 congruency effect for a fixated word by the congruency of a – lexically different – preview word would provide direct and convincing evidence for a parafoveal updating of sentence meaning by suggesting that sentence meaning immediately adapts to input from preview, which, in turn, affects the semantic integration of the fixated word. At a more general level, such evidence would support a generative and adaptive nature of the language processing system.

Eye tracking studies have shown that the congruency of the preview word can influence fixation times on subsequently fixated target words (Schotter, & Jia, 2016; Veldre & Andrews, 2016; Yang, Li, Wang, Slattery, & Rayner, 2014; Yang, Wang, Tong, & Rayner, 2012). However, in all of these studies, the foveal word was always congruent with the sentence. For this reason, it remains unclear whether foveal congruency processing (congruent vs. incongruent) can be influenced by the congruency of the preview. For example, longer reading times on foveal words after an incongruent parafoveal preview may just reflect the difficulty of processing the incongruent preview spilling over into the following fixations.

Recent ERP studies using the RSVP-with-flankers paradigm (Barber, Doñamayor, Kutas, & Münte, 2010; Barber, van der Meij, & Kutas, 2013; Li, Niefind, Wang, Sommer & Dimigen, 2015; Payne, Stites, Federmeier, 2019; Stites, Payne, Federmeier, 2017; Zhang, Li, Wang, & Wang; 2015) have provided some evidence on the effect of preview congruency by manipulating the congruency of both the preview word and the foveal word. In this paradigm, sentences are presented as word triplets during steady fixation, with the preceding word in the sentence appearing to the left and the subsequent word appearing to the right of fixation. During sentence presentation, the word triplet is updated step-wise, such that each word appears in preview before it moves into foveal vision (see also Figure 2). A major finding from this paradigm is that sentence-incongruent words in the preview position elicit a more negative N400 than congruent words. As N400 amplitude indexes lexico-semantic processing (Kutas & Federmeier, 2011), this effect was taken as evidence that parafoveal information is incrementally integrated into the sentence representation. More interestingly, N400 congruency effects were observed when an incongruent target word appeared in the preview position, but were reduced or absent after the word moved into foveal vision in the next step (Barber et al. 2010; Payne et al., 2019; Stites et al., 2017). These findings may suggest that information gleaned through preview can indeed influence subsequent semantic processing in the fovea.

Crucially, however, a potential confound in these previous ERP studies limits the interpretation of the foveal N400 effects observed. This confound is explained in Figure 1. Specifically, in all previous studies, congruency of the preview word and congruency of the subsequent foveal word were not independently manipulated but confounded with word identity. That is, the parafoveal and foveal words had always been the same, both in terms of lexical identity and sentence congruency (in the example of Figure 1, this is the case for the parafoveal “cave” followed by a foveal “cave” as well as for the parafoveal “cup” followed by a foveal “cup”). Put differently, in these studies, the foveal N400 congruency effect was attenuated and eradicated after a given word had been processed already in the parafoveal position. Therefore, any effects of congruency on foveal N400 amplitude were confounded with the effects of (parafoveal) identity priming (Rayner et al., 1975), as explained in the following. For foveal presentations, it is well-established that word repetition strongly attenuates the N400 (Rugg, 1985). Similarly, in reading with eye movements, a strong N400 reduction is observed if the same word is fixated twice across an intervening saccade (Dimigen, Kliegl, & Sommer, 2012). Furthermore, the N400 tends to be reduced if the fixated word was previously seen in parafoveal vision (identity preview benefit, Dimigen et al., 2012). Importantly, such priming effects due to identity repetition are not confined to the semantic level, but also involve several earlier stages of word processing. Reduced N400 effects of foveal congruency observed in previous studies may therefore be due to a confound with an identity preview effect on the N400.

**Figure 1.**
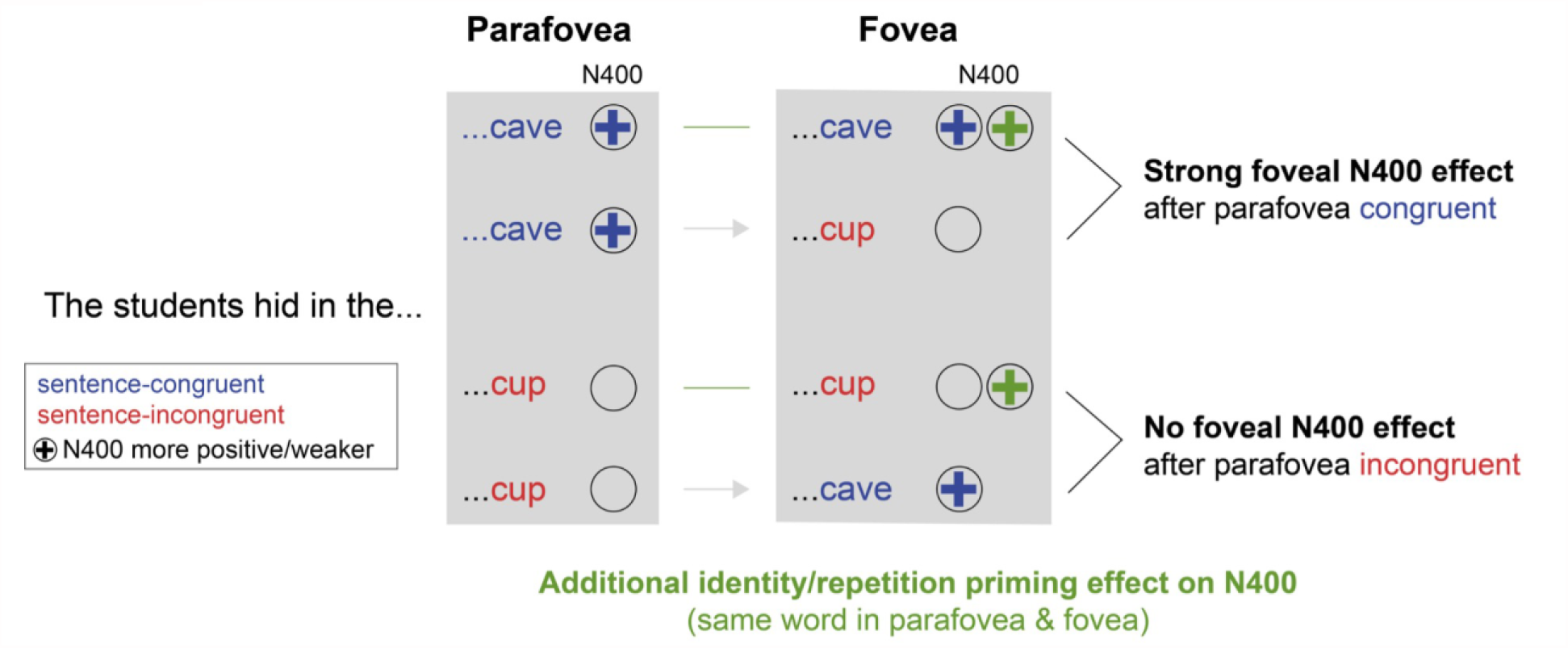
Illustration how foveal congruency effects can be confounded with word repetition effects (identity preview benefit) in the RSVP-with-flanker paradigm. In this figure, “plus” signs indicate a weaker (more positive) N400 amplitude. Previous studies have reported that the foveal N400 congruency effect is strong if the preceding parafoveal preview was sentence-congruent (marked in blue) but absent or attenuated if the preview was sentence-incongruent (marked in red). However, as illustrated here, this reported interaction between foveal and parafoveal congruency might also be simply explained by repetition priming effects (marked in green), which also would reduce the N400 amplitude in the two subconditions where the same word was shown both in the parafoveal and the foveal position. If this word repetition also leads to a weaker (more positive) N400, this effect alone could explain an observed interaction between parafoveal and foveal congruency.

In the present experiment, we build upon these earlier studies by manipulating whether the meaning of the preview and the subsequent foveal word are congruent with the sentence. Crucially, however, we also orthogonally manipulated the lexical identity between the preview and the foveal word, as being either identical or different words. When previewed and foveal word are identical, we should be able to replicate findings where the foveal congruency effect is diminished after the pre-processing of the (identical) preview. More importantly, this experimental design allows us to dissociate the foveal effect of preview congruency from the identity preview effect. Firstly, the new condition in which the preview and the subsequently fixated word were lexically different (but both congruent or both incongruent with the sentence), eliminates all non-semantic preview benefits. If preview congruency can still modulate the following foveal congruency effect under these conditions if the words are lexically different − this would clearly suggest that the semantic integration of the foveal word depends on the updating of sentential context through the preceding parafoveal preview, in line with generative theories of sentence processing.

## METHODS

### Participants

Of 31 right-handed native Chinese speakers with normal or corrected-to-normal visual acuity, volunteering for the experiment, one data set was removed due to a high rate of trial loss (70%). Of the remaining 30 participants (mean age = 23, range 21-27, years) 15 were women. This study was approved by the Psychology Research Ethics Committee of South China Normal University and participants provided written informed consent prior to the experiment.

### Materials

A total of 240 Chinese sentences were constructed, with target words embedded at the middle position of the sentence. The target was always a one-character word, but the remaining words in the sentence could consist of one, two, or three characters. In a 2 × 3 design, we orthogonally manipulated the sentence-congruency of the preview word, the sentence-congruency of the word later seen in the fovea, as well as the lexical identity of the preview word and the fovea word (same word or different words). This resulted in six conditions:

1. **preview congruent, foveal congruent (identical words)**
2. **preview congruent, foveal congruent (different words)**
3. **preview congruent, foveal incongruent**
4. **preview incongruent, foveal incongruent (identical words)**
5. **preview incongruent, foveal incongruent (different words)**
6. **preview incongruent, foveal congruent**

This design allowed us to answer four questions with regard to the influences on the N400 component during reading. First, we aimed to replicate findings from previous experiments where the foveal word’s N400 congruency effect vanished if the target word had already been seen in the parafovea (Barber et al. 2010; Payne et al., 2019; Stites et al., 2017). This was done by comparing the N400 in condition 1 (both words congruent and identical preview) with condition 4 (both words incongruent and identical preview). Second, the main effect of *parafoveal* congruency on the parafoveally-triggered N400 was tested by comparing the N400 in all congruent preview conditions (1, 2 and 3) with the N400 in all incongruent preview conditions (4, 5, 6). Third, the dependency of the *foveal* congruency effect on preview congruency – while controlling for identity preview effects – was tested by comparing the N400 in conditions 3 versus 2 – in which the preview word was congruent – with conditions (5) versus (6), in which the preview word was incongruent. Fourth, a possible identity preview/repetition priming effect on the N400 was isolated by comparing the conditions with an identical-word preview (1 and 4) to the conditions with a different-word preview (2 and 5). Because previous studies have reported identity preview effects on the N1 component, we conducted the latter analysis not only for the N400, but also for the N1.

Sixteen undergraduate students who did not participate in the main experiment rated the congruency of the preview words and foveal words with the sentences on a 5-point scale (1 = highly incongruent; 5 = highly congruent). Mean congruency ratings for congruent and incongruent preview words were 4.34 (*SD* = 0.61) and 1.72 (*SD* = 0.60), respectively; for the foveal words the corresponding congruency ratings were *M* = 4.40 (*SD* = 0.55) and 1.79 (*SD* = 0.57). As expected, congruency had a main effect on the rating, *b* = 2.65, SE = 0.04, *t* = 72.7, and there was no significant interaction with target word position (preview versus foveal), *b* = 0.03, SE = 0.04, *t* = 0.9. In the experiment, every target word that appeared as a congruent word in one sentence appeared as an incongruent word in another sentence, that is, lexical properties were strictly matched between congruency conditions.

Sentential constraint of the sentences was assessed in a cloze procedure study with 30 further undergraduate students from South China Normal University. Participants were presented with each sentence without the target word and were to complete the sentence with the first word that came to their mind. At the target word position, the mean cloze probability was 42% (*SD* = 20%).

### Procedure

Six sets of stimulus materials were created, each containing 240 sentences, with 60 sentences per condition. In the experiment, sentences were counterbalanced, such that each sentence frame was only shown once to each participant. After giving instructions and obtaining written informed consent, participants were assigned to one of six stimulus sets.

The trial scheme is shown in Figure 1 and closely resembles the one used by Li et al. (2015).

Each trial began with a fixation check of the eye tracker (see below), followed by the sentence presentation. Stimuli were displayed on a 17-inch CRT monitor (resolution 1024×768, vertical refresh rate 150 Hz). Sentences were presented as a series of triads of Chinese characters, displayed for 250 ms each and separated by 30-ms blank intervals, approximating a natural reading rate. Sentences were presented character-by-character at the centre of the screen, flanked by the preceding character to the left and the subsequent character to the right. Each character subtended 1.62°×1.62° degrees of visual angle. Characters were separated by one empty character space. Thus, the left edge of the right parafoveal flanker character was presented at an eccentricity of 2.43° (1.5 character spaces × 1.62°) from the screen center. Participants were to keep their gaze fixated on the central character, which was monitored by the eye tracker. After each sentence, participants were prompted to decide whether or not the sentence was semantically plausible by pressing the left or right mouse button.

### Recordings and data analysis

Eye movements of the right eye were recorded with an SR Eyelink 1000 eye-tracker (remote configuration) at 1000 Hz with head position stabilized via chin rest. Central fixation was verified gaze-contingently at the onset of each trial. The EEG was recorded from 42 Ag/AgCl electrodes mounted in a textile cap at standard positions of the 10-10 system. The electrooculogram was recorded from four electrodes positioned on the infraorbital ridge and outer canthus of each eye.

Online reference was the left mastoid, FCz served as ground. Electrode impedances were kept below 5 kΩ. Signals were amplified with Brain Products amplifiers at a time constant of 10 s and sampled at 500 Hz. Gaze and EEG were synchronized using the EYE-EEG extension (Dimigen, Sommer, Hohlfeld, Jacobs, & Kliegl, 2011) for EEGLAB (Delorme & Makeig, 2004). Offline, the EEG was high-pass filtered at 0.1 Hz (5^th^ order Butterworth filter) and low-pass filtered at 45 Hz (EEGLAB’s windowed sinc FIR filter with default settings). Finally, all channels were recalculated to average reference.

Trials with incorrect manual responses, EEG artifacts, or gaze samples outside the central fixation area during the presentation of the target screen were excluded (overall, 29% of trials). EEG epochs of accepted trials were segmented around the onset of the parafoveal target word, from −200 to 1,100 ms, and baseline-corrected by subtracting a 100-ms pre-stimulus baseline. To study N400 effects, we defined a ROI of four centroparietal electrodes (Cz, Pz, CP1, CP2) and two time windows from 300-500 ms after the onsets of the targets in the preview position and in the foveal position, respectively. In addition, to study the possible effects of word repetition (identity preview) on the N1 component reported in previous studies (e.g., Dimigen et al., 2012; Li et al., 2015), a second ROI of four occipitotemporal electrodes (PO9, PO7, PO8, PO10) was defined and analyzed for the time window from 200-300 ms after the onsets of the targets in the foveal position.

Single-trial EEG amplitudes were quantified as the average amplitude across the electrodes in the ROI and for each time window. Statistical analyses were performed using linear mixed effect models (LMM) with two fixed factors: (1) *preview congruency* (preview congruent or incongruent) and (2) *foveal congruency* (foveal word identical, congruent, or incongruent). Participants and items were included as crossed random factors, and a maximal random effects structure for intercepts and fixed factors was fit across participants and items.

## RESULTS

On average, participants responded correctly to the plausibility question after 87% of the trials. Response accuracy was significantly higher in the foveal incongruent condition (89%) than in the foveal congruent condition (83%), *b* = 0.52, SE = 0.09, *z* = 6.0, *p* < .001.

In the following, we will report the results with respect to the four possible effects on the N400 outlined in the Introduction:

### Question 1: Does the foveal N400 congruency effect vanish after previewing the same word?

We replicated the findings of previous experiments where the foveal word’s congruency effect vanished if the same word had already been preprocessed in the parafovea (Barber et al. 2010; Payne et al., 2019; Stites et al., 2017). By comparing the N400 in Condition 1 (preview congruent, foveal congruent, identical words), with Condition 4 (preview incongruent, foveal incongruent, identical words; see Table 1), we found the foveal word’s congruency effect to be diminished after parafoveal preprocessing (Figure 3). Specifically, the N400 congruency effect at the centroparietal ROI was marginally significant when congruent versus incongruent target words were shown in the parafovea, *b* = 0.47, SE = 0.25, *t* = 1.9, *p* = .06, but not when the same words were subsequently presented again in the fovea, *b* = 0.36, SE = 0.28, *t* = 1.3, *p* > .1. Interestingly, as evident from Figure 2, over more anterior brain regions there seemed to be an effect of foveal congruency and also of parafoveal congruency. Indeed, an exploratory post-hoc test at a frontal ROI (consisting of FPz, AFz, Fz) did reveal both a significant foveal congruency effect, *b* = 1.13, SE = 0.34, *t* = 3.3, *p* < .001, and a parafoveal congruency effect, *b* = 0.78, SE = 0.29, *t* = 2.6, *p* < .01.

**Table 1.** Example material and conditions

| Condition |  | Sentence | Congruency 1 | Congruency 2 |
| --- | --- | --- | --- | --- |
| preview congruent | (1) foveal congruent & identical | 同学们躲进山上的那个①洞-②洞来躲避风雨。 | 4.34 (0.61) | 4.34 (0.61) |
|  |  | The students hid in the ①cave-②cave in the mountain to avoid wind and rain. |  |  |
|  | (2) foveal congruent & different | 同学们躲进山上的那个①洞-②亭来躲避风雨。 |  | 4.40 (0.55) |
|  |  | The students hid in the ①cave-②pavilion in the mountain to avoid wind and rain. |  |  |
|  | (3) foveal incongruent | 同学们躲进山上的那个①洞-②杯来躲避风雨。 |  | 1.79 (0.57) |
|  |  | The students hid in the ①cave-②cup in the mountain to avoid wind and rain. |  |  |
| preview incongruent | (4) foveal incongruent & identical | 同学们躲进山上的那个①蛋-②蛋来躲避风雨。 | 1.72 (0.60) | 1.72 (0.60) |
|  |  | The students hid in the ①egg-②egg in the mountain to avoid wind and rain. |  |  |
|  | (5) foveal incongruent & different | 同学们躲进山上的那个①蛋-②亭来躲避风雨。 |  | 4.40 (0.55) |
|  |  | The students hid in the ①egg-②pavilion in the mountain to avoid wind and rain. |  |  |
|  | (6) foveal congruent | 同学们躲进山上的那个①蛋-②杯来躲避风雨。 |  | 1.79 (0.57) |
|  |  | The students hid in the ①egg-②cup in the mountain to avoid wind and rain. |  |  |

*Note:* The preview for the target word was either congruent (here: *cave*) or incongruent (here: *egg*) with the sentence context, and the later foveal target word was presented either in its identical (here: *cave or egg*), congruent (here: *pavilion*) or incongruent version (here: *cup*), yielding a $2 \times 3$ design (*preview congruency* × *foveal congruency*). The target always consisted of a single Chinese character. ① Target in the preview position. ② Target in the later foveal position. “Congruency 1” refers to the plausibility rating of how well the preview word fit into the first part of the sentence (up to and including the target word). “Congruency 2” refers to the plausibility rating of how well the foveal word fit into the first part of the sentence. For both ratings, 1 is “highly incongruent”, 5 is “highly congruent”.

**Figure 2.**
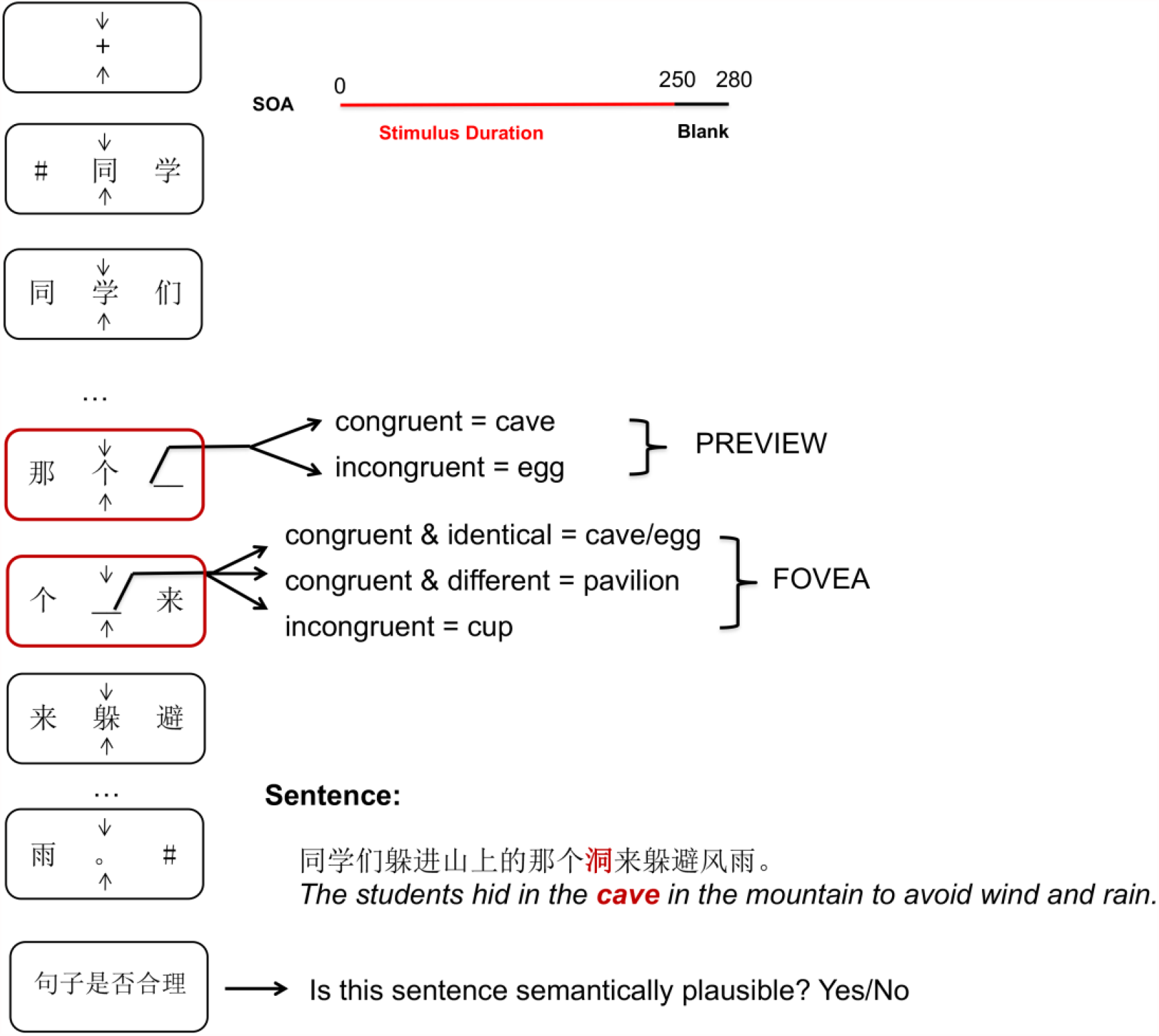
Trial scheme for the experiment. Sentences were presented character-by-character in the center of the screen, flanked by the preceding character to the left and the subsequent character to the right. The target word at the parafoveal position was presented either in its congruent (here: cave) or its incongruent version (here: egg). The target word in the foveal position was presented either in its identical (here: cave/egg), congruent (here: pavilion) or incongruent version (here: cup). Character triads were displayed for 250 ms each and separated by 30 ms blank intervals. Participants were asked to keep their gaze fixated on the central character (monitored with eye-tracking) and to judge the plausibility of the sentence with a button press at the end of the trial.

**Figure 3.**
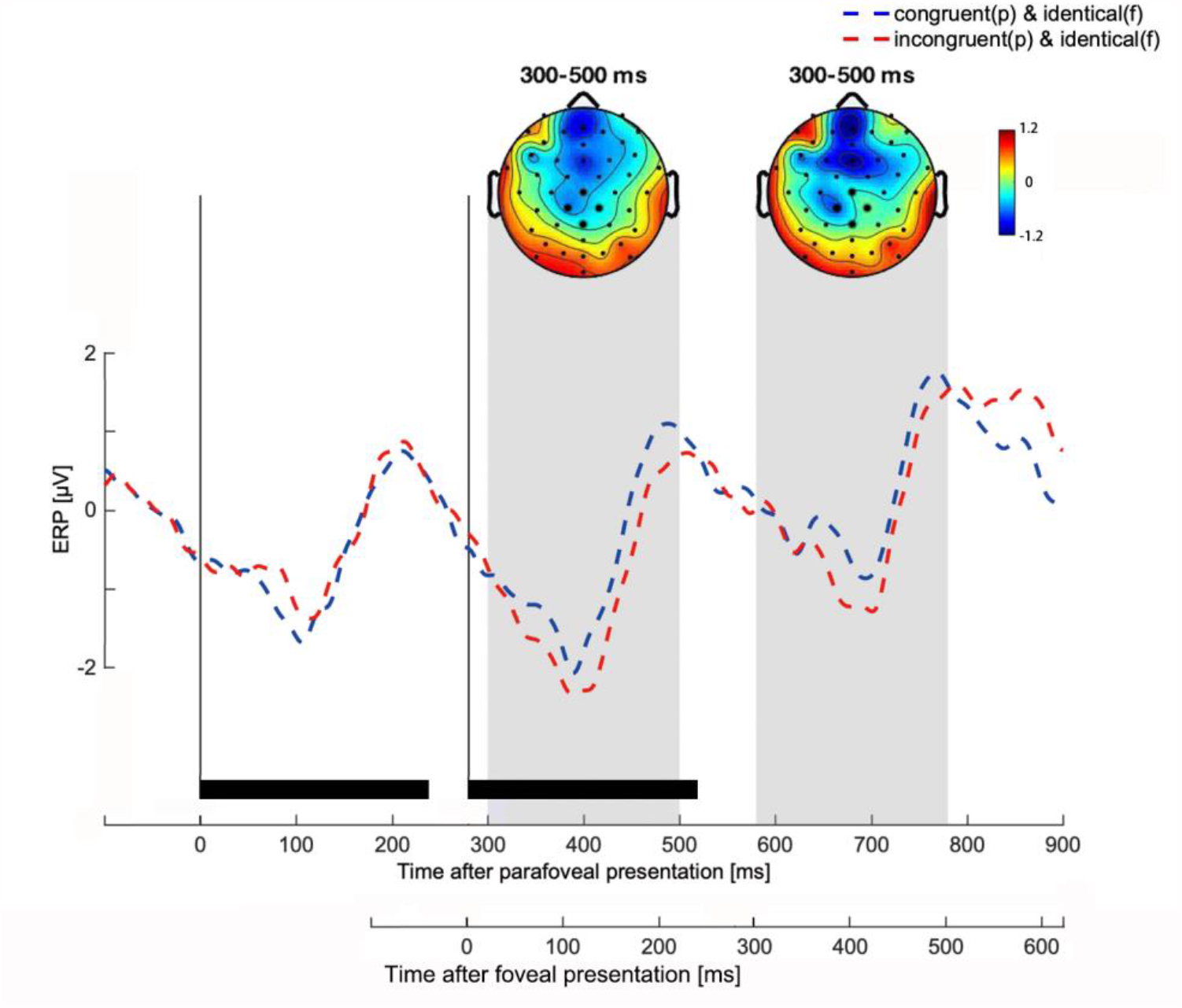
ERP congruency effects for the parieto-central ROI when a target word appeared in the parafovea and subsequently again in the fovea. Time 0 on the time axis marks the onset of the target word in the right parafoveal position. Black bars above the time axis mark the stimulus durations of the character triads. The two vertical lines mark the onset of the triad containing the target word in the parafovea and fovea, respectively. ERP waveforms show the average activity in the ROI of four centroparietal electrodes (highlighted by larger black markers in the topographic maps) for the congruent and incongruent conditions. Scalp topographies show the difference between condition 4 (preview incongruent, foveal incongruent & identical) minus condition 1 (preview congruent, foveal congruent & identical condition) from 300-500 ms after target onset in the parafoveal and fovea, respectively (see grey shaded areas).

### Question 2: Can we replicate a parafoveal N400 congruency effect?

Previous RSVP-with-flanker studies have reported an N400 congruency effect following the presentation of the word already in the parafoveal position (Barber, et al., 2010; Barber, et al., 2013; Li, et al., 2015; Stites, et al., 2017; Payne, et al., 2019; Zhang, et al., 2015). Here, we aimed to replicate this effect by comparing all congruent parafoveal conditions (conditions 1,2 and 3) with all incongruent parafoveal conditions (conditions 4, 5 and 6) in the window between 300–500 ms after target onset in the *parafovea*. Indeed, incongruent parafoveal targets elicited a significantly more negative N400 than congruent parafoveal targets over centroparietal scalp sites, *b* = 0.44, SE = 0.16, *t* = 2.7, p <.01. This suggests semantic integration occurring while words are still in the parafovea (see Figure 4).

**Figure 4.**
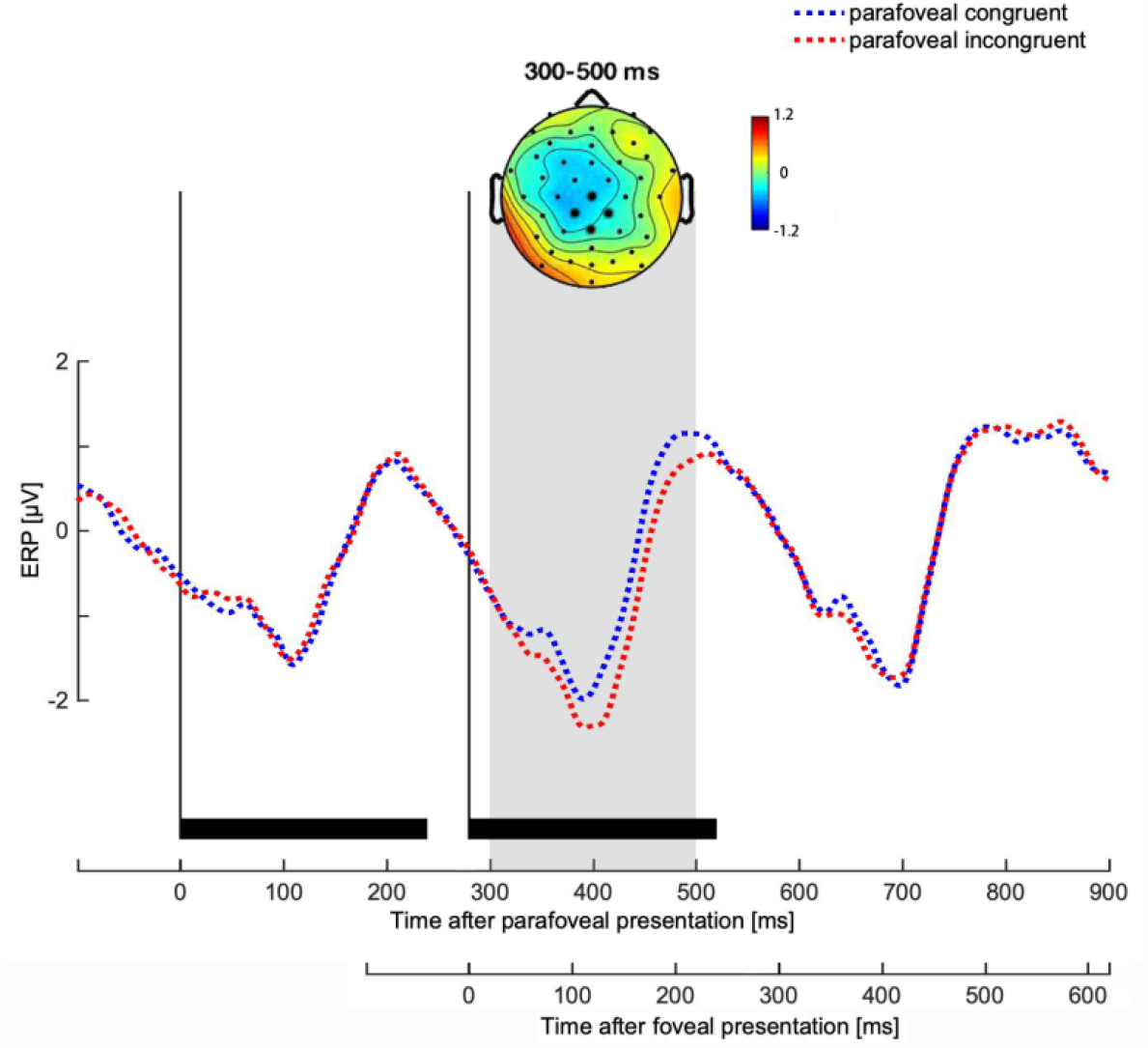
Effect of parafoveal congruency on the N400 component. Waveforms show the average activity in the ROI of four centroparietal electrodes (highlighted in black in the topographic map) for the parafoveal congruent and parafoveal incongruent conditions. The topographic map shows the difference in ERP activity (incongruent parafoveal conditions minus congruent parafoveal conditions) in the time window of 300-500 ms after the onset of the target word in the parafoveal position.

### Question 3: Does the foveal congruency effect depend on preview congruency?

As a novel aspect of our study, we tested the dependency of the *foveal* congruency effect on preview congruency in the conditions in which *different* words were seen in both positions. That is, we compared the N400 in conditions (3) versus (2) – where the previewed words were congruent – with conditions (5) versus (6) – where the previewed words were incongruent (Figure 5, middle). We observed a significant interaction between the congruency of the foveal word with the congruency of the preceding parafoveal preview, *b* = 0.81, SE = 0.40, *t* = 2.0, *p* <.05. Specifically, when preview had been a congruent (different) word, we found a significant N400 effect of semantic congruency during the subsequent foveal target presentation (*preview congruent, foveal incongruent condition versus preview congruent, foveal congruent & different condition;* conditions 3 vs. 2), *b* = 1.01, SE = 0.28, *t* = 3.5, *p* < .001. In contrast, when the preview word had been an incongruent (different) word, there was no indication for a foveal congruency effect on the N400 (*preview incongruent, foveal incongruent & different condition versus preview incongruent, foveal congruent condition;* conditions 5 vs. 6), *b* = 0.19, SE = 0.28, *t* = 0.7, *p* > .4. A follow-up pair-wise comparison between the *preview congruent, foveal incongruent* and the *preview incongruent, foveal incongruent* & different conditions (conditions 3 vs. 5) showed that incongruent foveal words elicited a more negative N400 following a congruent as compared to an incongruent preview, *b* = 1.13, SE = 0.30, *t* = 3.8, *p* < .001. Taken together, these results suggest that the semantic integration of a word in the fovea is modulated by the congruency of the word in preview, even under strict control of (parafoveal) identity priming effects on the N400.

**Figure 5.**
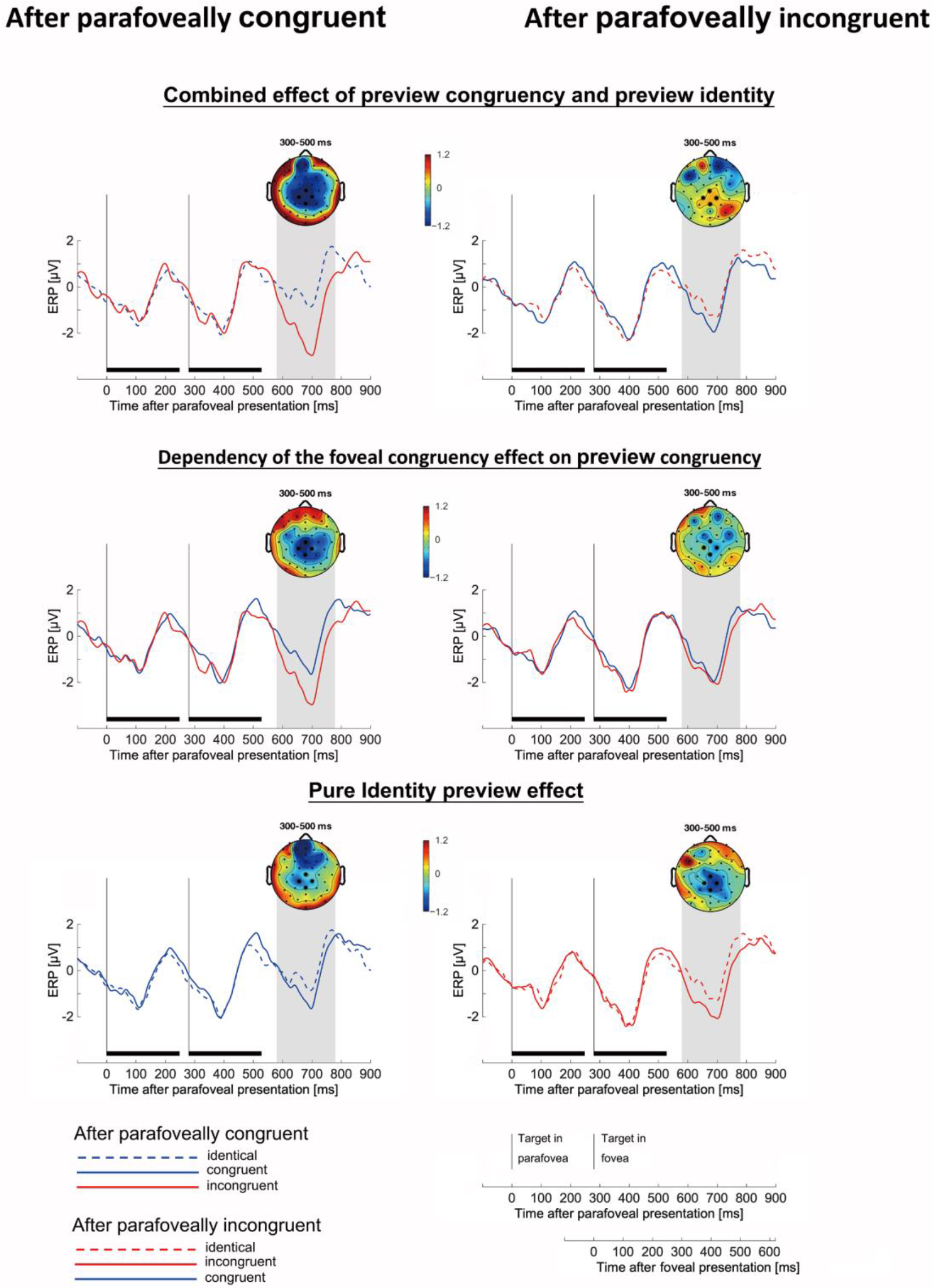
Disentangling the influences on the foveal N400 effect and its interaction with the preview word. To separate congruency and identity preview effects, the six conditions can be plotted in three ways. Top. Combined (and confounded) effects of preview congruency and preview identity. Middle. Dependency of the foveal congruency effect on preview congruency under control of identity preview effects. Bottom. Pure identity preview effect. Left: N400 after parafoveally congruent word; Right: N400 after parafoveally incongruent word.

To compare how the parafoveal and foveal congruency interaction looks with and without the control for preview identity effect, in addition to the preview congruency effect and identity preview effect, we tested as a supplementary analysis, the combination of the preview congruency effects and preview identity effects (Figure 5, top). The combinatory effects were tested as the dependency of the *foveal* congruency effect on preview congruency in the conditions in which the same words were seen in both positions if they were same in the sentence congruency, that is comparing the N400 in conditions (3) versus (1) – in which the previewed words were congruent – with conditions (4) versus (6), in which the previewed words were incongruent. When these conditions were analyzed, there was a significant interaction between the congruency (and with identity) of the preview word and the congruency of the foveal word, *b* = 0.95, SE = 0.39, *t* = 2.4, *p* < .05. Specifically, when the preview had been congruent (and half was identical with the target word), we found a significant N400 effect of semantic congruency during the subsequent foveal target presentation (*preview congruent, foveal incongruent* condition versus *preview congruent, foveal congruent & identical* condition; conditions 3 vs. 1, *b* = 1.48, SE = 0.30, *t* = 4.9, *p* < .001). In contrast, when the preview word had been incongruent (and half was identical with the target word), there was no indication for a foveal congruency effect on the N400 (*preview incongruent, foveal incongruent & identical* condition versus *preview incongruent, foveal congruent* condition; conditions 4 vs. 6, *b* = 0.52, SE = 0.30, *t* = 1.7, *p=*.*083*.

### Question 4: Is there an identity preview effect on the N400 (and N1) component?

Finally, we tested for any identity preview effects in the N400 time window by comparing conditions with identical preview (1 and 4) to conditions with different-word previews (2 and 5), while controlling for congruency. Results are shown in Figure 5 (bottom). Importantly, we found a main effect of identity preview in the N400 time window (*b* = 1.00, SE = 0.22, *t* = 4.4, *p* <.001). As to be expected from the repetition priming literature (see also Dimigen et al., 2012), invalid, different-word previews elicited a more negative N400 than valid, identical previews. This identity preview effect was independent of the congruency of the previewed word (*b* = 0.25, SE = 0.39, *t* = 0.6, *p* > .5).

The identity preview effect also appeared at the occipitotemporal ROI, 200–300 ms after foveal target word onset (Figure 6). Specifically, invalid previews elicited more negative N1 component amplitudes compared to valid, identical previews (*b* = 0.68, SE = 0.22, *t* = 2.9, p <.01), replicating the N1 effect (“preview positivity”) previously observed in both fixation-related potentials (Degno et al., 2019; Dimigen et al., 2012; Kornrumpf, Niefind, Sommer, Dimigen, 2016; Niefind & Dimigen, 2016) and ERPs (Li et al., 2015). Interestingly, we also observed an unexpected interaction between this early N1 identity preview effect and the congruence of the preview words (*b* = 0.91, SE = 0.39, *t* = 2.3, *p* < .05). Specifically, the N1 effect only appeared when the previewed word and the foveal word were both incongruent with the sentence context (*preview incongruent, foveal incongruent & different* condition versus *preview incongruent, foveal incongruent & identical* condition; conditions 5 vs. 4), *b* = 1.01, SE = 0.32, *t* = 3.2, p < .01, but not when they were both congruent with the sentence (*preview congruent, foveal congruent & different* condition versus *preview congruent, foveal congruent & identical* condition; conditions 2 vs. 1), *b* = 0.10, SE = 0.32, *t* = 0.3, *p* > .7.

**Figure 6.**
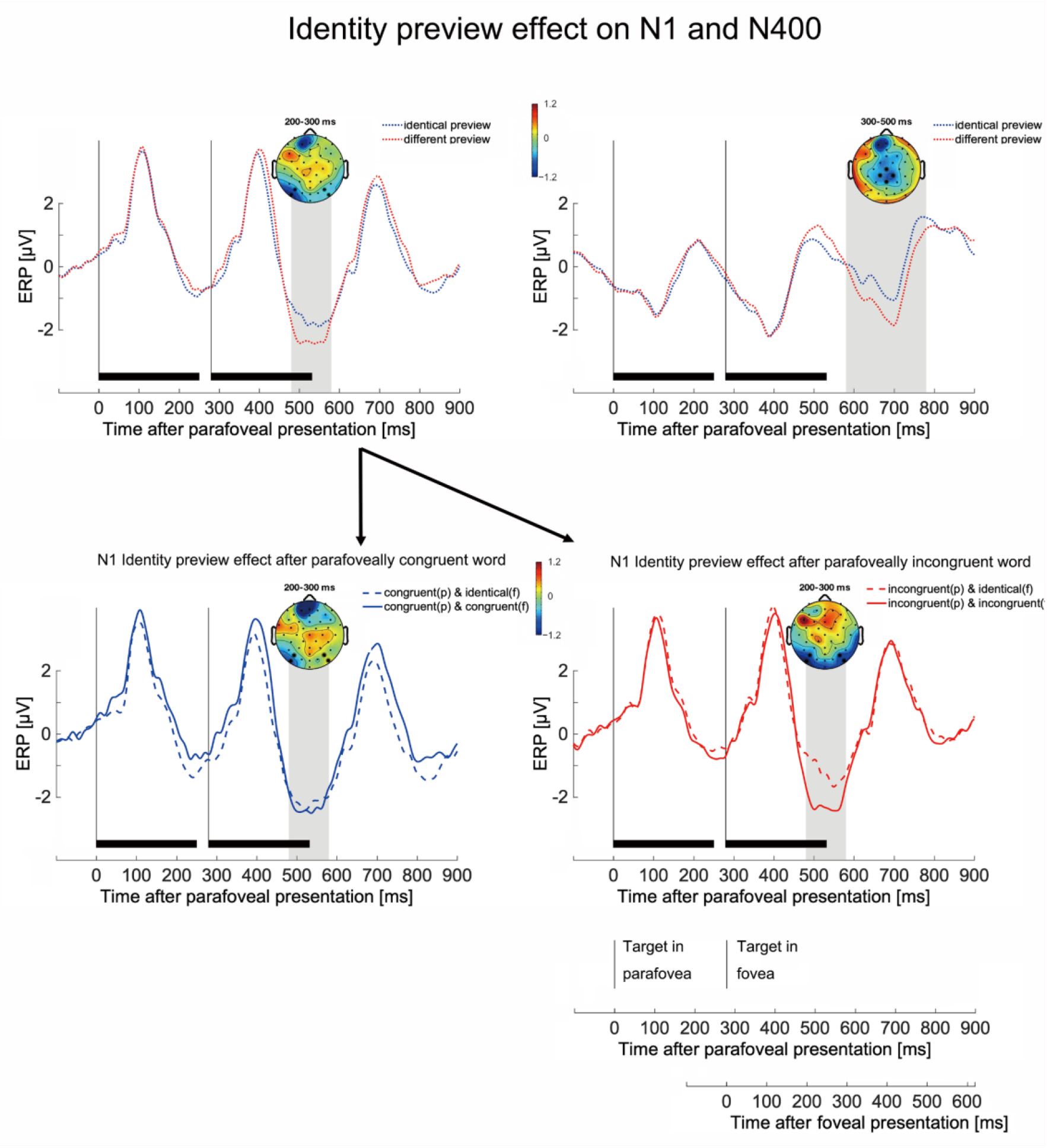
Top: Main effect of preview identity on the early N1 component at occipitotemporal electrodes (Left) and on the later N400 component at the centroparietal ROI (Right). Bottom: Unexpectedly, the early N1 preview validity effect (at occipitotemporal electrodes) only appeared when the preview word was sentence-incongruent (preview incongruent, foveal incongruent & different condition versus preview incongruent, foveal incongruent & identical condition; conditions 5 vs. 4; right panel) but not when the preview word was sentence-congruent (preview congruent, foveal congruent & different condition versus preview congruent, foveal congruent & identical condition; conditions 2 vs. 1; left panel). Waveforms show the average activity in the ROI of four electrodes (highlighted in black in the topographic map) for the identical preview condition and different preview condition. Topographic map shows the difference in ERP activity (different preview conditions minus identical preview conditions) in the time window of 200-300 ms (Top-left and Bottom) and 300-500 ms (Top-right) after the onset of the word in the foveal position.

## DISCUSSION

In a RSVP-with-flankers task and using the N400 component of the ERP, we investigated whether the sentential information is rapidly updated based on parafoveal information, thereby already changing the context for word when it is fixated word. To address this question, we orthogonally manipulated the sentence congruency of both the preview word and the subsequently fixated words during reading. In addition, we also manipulated the lexical identity between the previewed word and the later foveal word – as being either the same or different words – in order to distinguish genuine effects of contextual processing from any non-semantic effects of parafoveal-to-foveal repetition priming (i.e., an identity preview effect).

First, we replicated the finding of previous studies that the N400 effect of foveal word’s congruency is diminished after preprocessing the same word in preview. Second, we replicated the N400 effect elicited by the congruency of parafoveally presented words reported in previous studies. Third, and more importantly, we observed a clear interaction between the sentence congruency of the preview word and that of the subsequently fixated foveal word in predicting N400 amplitude.

Importantly, this was even the case if the previewed word and the foveally presented word were lexically different, excluding any effects of repetition priming on the N400 (which we also isolated, as discussed below). Specifically, the N400 effect of foveal word congruency was only present when the word previously seen in preview had been congruent with the sentence context. These results suggest that readers utilize the sentence context generated from parafoveal information to modulate the subsequent foveal processing of words, and that this preview effect occurs at the level of semantic integration. To our knowledge, this is the first study direct evidence about a preview effect on subsequent foveal processing at the semantic integration level. Fourth, if the same word was visible in the parafovea and in the fovea, we found a significant identity preview effect not only on the early N1 component (e.g., Dimigen et al. 2012), but also on the later N400 component. To our knowledge, this is the first clear demonstration that an identical parafoveal preview – which is of course the norm in natural reading – attenuates the N400 component once the word is directly fixated. In the following, we will discuss these findings in more detail.

### Question 1: Does the foveal N400 congruency effect vanish after previewing the same word?

Previous studies show that N400 congruency effects are reduced or absent after the parafoveal word moved into foveal vision (Barber et al. 2010; Payne et al., 2019; Stites et al., 2017). While our results show a reduction of the foveal congruency effect on the centroparietal N400 component after seeing the same word in the parafovea, they also suggest that the foveal congruency effect is shifted towards anterior regions, which is also consistent with the data patterns reported by Stites et al. (2017; their Figure 3) and in a study with Chinese readers (Zhang et al., 2015). This result indicates the involvement of cognitive processes reflected by anterior scalp distributions of N400 effects, such as situation establishment and thematic relationships (e.g., Chwilla & Kolk, 2015; Liang, Xiao, Lei, Li, & Chen, 2020), word familiarity (e.g. Curran, 2000; Paller, Voss, & Boehm, 2007) or conceptual priming (Voss & Federmeier, 2011), which may shift the typical parietal foveal N400 effect towards anterior regions when foveal words had been preprocessed in the parafovea.

### Question 2: Can we replicate a parafoveal N400 congruency effect?

Our finding of an N400 congruency effect following the presentation of the word in the parafoveal position is in line with previous studies that have reported such an effect in the RSVP-with-flankers paradigm (Barber, et al., 2010; Barber, et al., 2013; Li, et al., 2015; Payne, et al., 2019; Zhang, et al., 2015; Stites, et al., 2017). The finding that the congruency of a parafoveal word modulates the N400 in the RSVP-with-flanker paradigm suggests that readers can quickly integrate the meaning of a not-yet-fixated word into the sentence.

### Question 3: Does the foveal N400 congruency effect depend on preview congruency?

More importantly, we found that the semantic integration of words in the fovea is modulated by the semantic congruency of the previewed words. In the present study, a foveal N400 effect – reflecting semantic integration of the foveal word – appeared only when the – non-identical – parafoveal preview had been congruent with the sentence context, but not when it had been incongruent. This suggests that the processing of an incongruent preview word hampers the semantic integration of a subsequent – and also incongruent – word in the fovea.

We see several possible explanations for the apparent interference that the preview caused in this condition. First, the incongruent word in the preview position may have altered the sentence context in such a way that the semantic integration of the foveal word was not possible or – more likely, given the generally small negative amplitude of the incongruent foveal word elicited in the N400 time window – not attempted or by-passed. Second, the semantic integration of a word into an incongruent context is an effortful, resource-limited process. Hence, it may be difficult to perform semantic integrations twice within a short interval because the semantic system may be refractory after the first time. Alternatively, the semantic integration stage of word processing may be able to handle only one word at a time but may take longer than the interval between two triplet presentations in the present study. In this case, semantic integration would constitute a central processing bottleneck. Such resource limitations for semantic processes have been reported, for example, for the N400 elicited by word pairs by Hohlfeld, Sangals, and Sommer (2004). Although elucidating the precise reasons for the absence of semantic integration effects after incongruent parafoveal information must be subject to future research, the present results indicate that our semantic system has problems to complete two integration processes in short succession or within the same sentence.

More generally, our finding that the semantic integration of fixated foveal words adapts to the sentential meaning generated based on the parafoveal preview strongly supports a generative and adaptive view of language processing. Generative models of language comprehension such as the Bayesian model assumes that the behavior of reader follows entirely from the requirement to make optimal decisions based on the available information (Norris, 2006). From this perspective, any generative model of language comprehension must consider message-level representations as probabilistically generating information at multiple types and levels of representation (Kuperberg & Jaeger, 2015). The preview effect of sentential congruency in the present study demonstrates the rapid availability of information generated from message-level representations. That is, the sentential meaning updated through brief information from preview can still contribute to the meaning construction of later words in the fovea. Our findings therefore support a dynamic and actively generative framework of language comprehension, in which message-level representations play an immediate and pivotal role.

### Question 4: Is there an identity preview effect on the N400 (and N1) component?

In the current study, we independently manipulated congruency and identity, which allowed us to disentangle their effects and to examine both in isolation and in interaction. Our results show that the foveal N400 amplitude was modulated by both preview congruency and preview validity (identical word vs. different word). As a novel result, we found a clear identity preview effect not only on the earlier N1 component – replicating the findings of Dimigen et al. (2012), Li et al. (2015) and others – but also on the late N400 component, which had only showed a statistical trend in the same direction in a previous study (Dimigen et al., 2012). The early N1 effect of preview validity has previously been interpreted as a reflection of priming at the level of orthographic or phonological representations of the upcoming word (Dimigen et al., 2012). In contrast, the late effects of preview validity on the N400 may reflect a form of repetition priming at the level of semantics. We believe that the finding of a reduced N400 by preview is important, because during normal reading, readers always see a correct (identical) preview on the upcoming word. Our finding therefore suggest that the N400 may be routinely attenuated under natural reading conditions as compared to more artificial laboratory conditions that do not provide a normal preview on the next word (e.g., traditional single-word RSVP paradigms).

Unexpectedly, we also found that the earlier N1 identity preview effect was modulated by the congruency of the preview word. Specifically, the N1 preview identity effect was apparent only when the preview word could not be easily integrated into the current sentence context. This indicates that the message level information may override the lexical priming effects on orthographical or phonological levels. This unexpected but interesting interaction deserves further study research with dedicated experiments in the future.

## CONCLUSION

In our experiment, we dissociate identity priming from semantic effects of parafoveal words.

While a correct preview of the upcoming word generally attenuates the foveal N400 component, regardless of congruency, parafoveal preview of semantically incongruent words interferes with the semantic integration of foveal information. These results suggest that word processing rapidly adapts to the sentential meaning generated from parafoveal previews and supports the notion of a dynamic and actively generative language processing system, which promotes fast and effective reading.

## AUTHOR NOTE

This work was supported by the National Natural Science Foundation of China (grant: 31700992). The funders had no role in study design, data collection and analysis, decision to publish, or preparation of the manuscript. The authors declare that the research was conducted in the absence of any commercial or financial relationships that could be construed as a potential conflict of interest.

